# ZIPPI: proteome-scale sequence-based evaluation of protein-protein interaction models

**DOI:** 10.1101/2023.05.25.542344

**Authors:** Haiqing Zhao, Donald Petrey, Diana Murray, Barry Honig

**Affiliations:** Department of Systems Biology, Columbia University Irving Medical Center, New York, NY 10032, USA; Department of Biochemistry and Molecular Biophysics, Columbia University Irving Medical Center, New York, NY 10032, USA; Department of Medicine, Columbia University, New York, NY 10032, USA; Zuckerman Mind Brain and Behavior Institute, Columbia University, New York, NY 10027, USA

**Keywords:** Protein-protein interactions, protein sequence, coevolution

## Abstract

Predicting protein-protein interactions (PPI) is a challenging problem of central importance in fundamental biology. With the increasing number of available PPI prediction methods and databases, an effective evaluation model would be extremely valuable. Here we introduce ZIPPI (Z-score for Information about Protein-Protein Interfaces), which evaluates structural models of a complex based on sequence co-evolution and conservation involving residues that are in contact in the interface. The interface Z-score (ZIPPI score) is calculated by comparing metrics for interface contacts to metrics obtained from randomly chosen surface residues. Since contacting residues are defined by the structural model, this obviates the need of accounting for indirect interactions with methods such as Direct Coupling Analysis. Although ZIPPI relies on species-paired multiple sequence alignments, its focus on contacting interfacial residues and the avoidance of direct coupling methods makes it computationally efficient. The performance of ZIPPI is evaluated through applications to experimentally determined complexes from the Protein Data Bank (PDB) and to decoys from the Critical Assessment of PRedicted Interactions (CAPRI) experiment. We demonstrate how ZIPPI can be implemented on a genome-wide scale by calculating scores for millions of structural models of protein-protein interactions in the *E. coli* interactome as predicted by PrePPI. Many PrePPI predictions filtered by ZIPPI score are novel. In all, this proteome-scale method shows promising feasibility for applications to the full human protein interactome, which is not yet accessible to deep learning methods.

## Introduction

The past decade has seen continuing developments in the prediction of protein-protein interactions (PPIs). One can trace these advances to the use of amino acid coevolution to predict inter-residue contacts^1,2^. These methods have been used to predict the structures of small proteins^3–5^ and, more recently, to predict interaction partners and interfacial residues involved in PPIs^6–9^. The underlying premise is that functional interactions between two residues will result in their coevolution, which should be reflected in paired multiple sequence alignments (MSAs) of putative orthologues and detectable through mutual information (MI) based metrics between the two positions in the alignment. A serious complication is that the correlation between two residue positions ***i*** and ***j***, i.e., two columns in the MSA, may result from an indirect coupling of ***i*** and ***j*** through their interaction with a third residue ***k***. To solve this problem, methods using Potts model^10^ including Direct Coupling Analysis (DCA)^3,11,12^ and EVcouplings^6,8^, sparse inverse covariance (PSICOV)^13,^ or pseudolikelihood maximization method such as Gremlin^4,7^ have been developed. However, these methods rely on the availability of large MSAs and thus have almost exclusively been applied to bacterial systems. Our main focus in this work is *E. coli* as well but, as will be discussed, there is no barrier to applications to larger genomes.

The advent of AlphaFold^14^ and other deep learning-based methods^15,16^ has fundamentally changed the landscape of sequence-based PPI prediction. These methods depend either directly on MSAs or may learn them from training on a large number of sequences using various deep language models^17–20^. Some methods predict contacting residues in PPIs while others predict the structures of multimeric complexes by concatenating the sequences of two proteins and folding them together as if they were a single protein (see e.g. AlphaFold-Multimer^21^ and AF2Complex^22^). An underlying problem for MSA-based methods is that, for a heterodimeric pair, it is necessary to carry out a species-based matching of the two query sequences which limits application to eukaryotic organisms due to the relatively limited number of sequences available for a paired MSA. Baker and coworkers discussed the challenges associated with applying deep learning and co-evolution-based methods to eukaryotes and were able to apply a hybrid RoseTTAFold/AlphaFold approach to a portion of the yeast proteome enabled in part by the large number of fungal genomes available^23^. However, applying deep learning to predict whether and how two proteins interact for entire proteomes remains computationally prohibitive and is likely to remain a significant challenge for some time.

The focus on sequence has, in a sense, diverted attention from the use of three-dimensional structure information to predict whether and how two proteins interact. Docking-based methods predict models of dimers based on the structures of interacting monomers^24,25^ but have not been applied on a proteome-wide scale and have not been used on this scale to predict *whether* two proteins interact. Template-based modeling^26^ is an alternate approach where the structures of individual proteins are superimposed on structurally similar proteins that appear in a complex present in the PDB^27^. In a series of papers we reported the PrePPI (Predicting Protein-Protein Interactions) algorithm^28^ and database^29–31^ that relies on template-based modeling and, through a novel and highly efficient scoring function, leverages structural information on a truly proteome-wide scale. For example, PrePPI effectively screens the ∼200 million possible pairwise combinations of human proteins which, in practice, amounts to billions of possible domain-domain interactions.

Here we use PrePPI predicted complex structures in *E. coli* to examine the extent to which simple evolution-based metrics are informative even in those cases for which the multiple sequence alignment (MSA) depth is shallow. Co-evolution methods are based on sequence alone without prior knowledge of interfacial residues. We turn this problem around and ask if, given an interface (predicted for example by PrePPI), can one use covariance across the interface to discriminate cognate from non-cognate binding partners? This problem is much simpler than the more general one which begins with sequence, and requires methods such as DCA, since it involves evaluating interfaces where interacting residues are already defined. Thus, we expect that MI calculations alone would be sufficient, even for eukaryotic proteins, as very deep MSAs required for DCA analysis would not be necessary. Our method, ZIPPI (for Z-score Information for Protein-Protein Interfaces), uses MSAs to determine coevolutionary information across interfaces but also leverages sequence conservation which provides an additional signal as to the reliability of a predicted interface. An essential feature of ZIPPI is the comparison of predicted interfaces evaluated with evolutionary metrics derived from MSAs to those created by replacing interfacial residues with randomly chosen surface residues.

Our focus on interfacial residues leads to a significant speedup in interface evaluation that allows us to apply ZIPPI on a genome-wide scale. Similarly, our finding that DCA is not needed for heterodimeric complexes effectively removes the need for large species-paired MSAs. As shown below ZIPPI is extremely effective in distinguishing correct from incorrect protein-protein interfaces as indicated by tests on PDB structures and on a CAPRI benchmark set^32,33^. Most notably there is a strong inverse correlation between ZIPPI scores and false positive rates (FPRs) for PrePPI predictions thus providing strong support for the reliability of ZIPPI’s efficacy and applicability to genome-wide interactomes.

## Results

### ZIPPI overview

Given a structural model of a protein-protein complex, we first identify interfacial contacts and all surface residues. A species-paired MSA is created and used to calculate the following metrics for the interfacial contacts: mutual information (MI), conservation (Con) and direct coupling (DCA). For each of these, an average product correction (APC)^34^ is used to remove the random and background signal potentially arising from an insufficient number of sequences in the MSA and the common phylogenetic relationships of the species represented in the alignment. We then calculate the average value of each metric for the entire interface and, in addition, retain the highest scoring contact, denoted as “top”, for each of the six metrics, resulting in a total of twelve metrics that characterize a predicted interface.

The next step is to substitute the contacting residues in the interface with a set of randomly chosen surface residues that are not in the interface (Figure 1). A hundred such interfaces are generated in this way for each complex. A Z-score of the predicted interface is then calculated for each of the twelve metrics based on a comparison to the corresponding values for the randomly generated interfaces.

**Figure 1.**
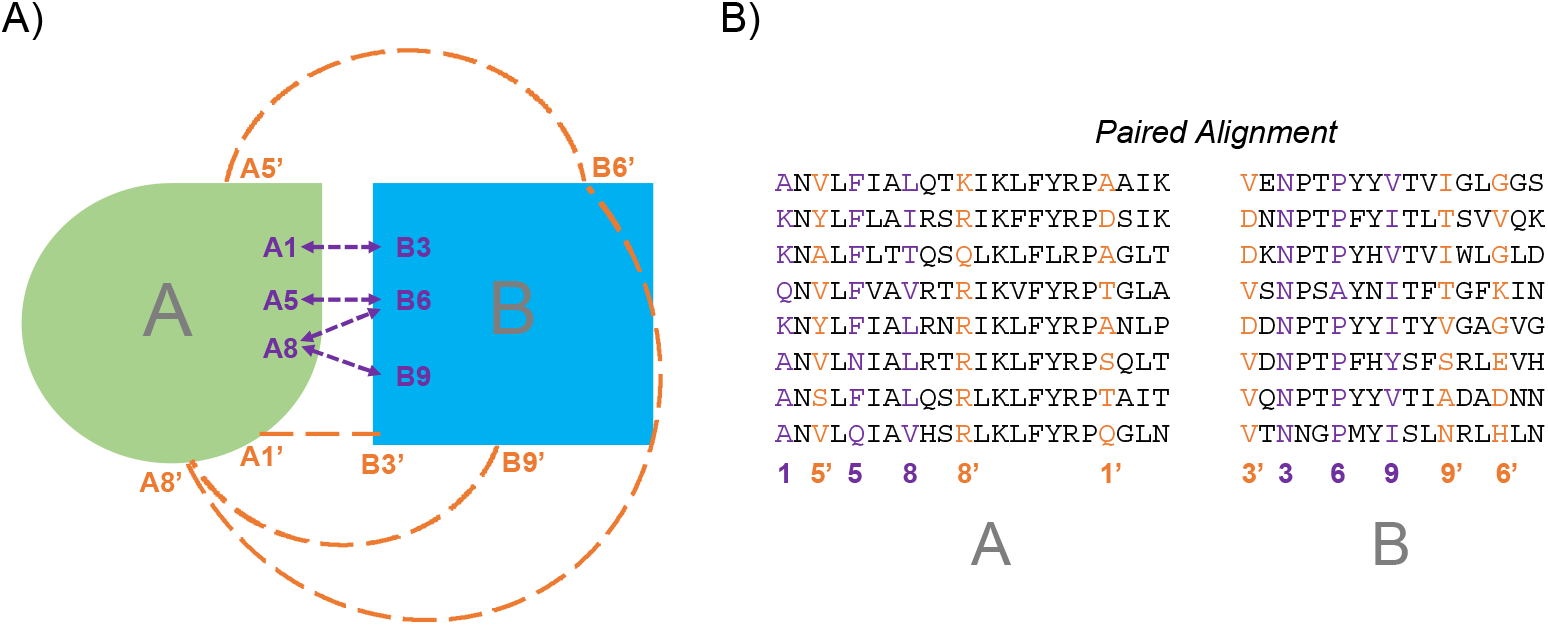
Schematic of the ZIPPI algorithm. Two proteins, A and B, form a complex with interfacial contacts between residues A1 and B3, A5 and B6, A8 and B6 or B9. Various evolutionary metrics (see text), calculated from the paired multiple sequence alignment in the right panel, are calculated for these contacting residues. The columns in the alignment that correspond to these interface residues are marked in purple. The false interfaces are generated by replacing the interface residues with randomly picked surface residues that are not on the interface. The dashed orange lines in the left panel indicate one such interface and the corresponding columns in the alignment are marked in orange.

### Testing ZIPPI on PDB complexes

Dimeric PDB complexes were collected from the first bioassembly as defined in the PDB structure file, for both bacterial and human complexes allowing us to test performance on both prokaryotic and eukaryotic MSAs with very different MSA depths. As described in Methods, complexes were selected based on resolution, chain length and the requirement that the proteins from the same species are present in the complex. In total, we obtained results for 279 bacterial heterodimers, 247 human heterodimers, 3976 bacterial homodimers and 717 human homodimers. For each complex we calculated values of the 12 metrics.

Figure S1, for each of the twelve metrics, plots the fraction of PPIs with a Z-score above the threshold denoted along the x-axis. It is evident that, for bacterial and human heterodimers, the APC correction improves performance relative to raw (uncorrected) metrics for MI and DCA but not for Con. In contrast, for homodimers the APC corrected Con metric is more effective than the corresponding raw metric. Further, choosing the top value for each metric is less effective than choosing the value averaged over the entire interface (Figures S1-S2, Figure 2). This is not unexpected since all contacts identified in PDB complexes are presumed to be correct and likely contribute to the total score. In Figure 2, we have plotted a single curve for the average value of MI, Con, DCA and their “top” equivalents. For each, the metric that contributes the highest Z-score is chosen (e.g. the MI curve is based on choosing the larger of the Z-values from <MI> or <MI_apc_>). In addition, the ZIPPI curve is generated by choosing the maximum value over all twelve metrics for each interface. In the following, for any property, the ZIPPI score will always correspond to the maximum value of all metrics such that the score for one interface may be based on a different metric than for another interface.

**Figure 2.**
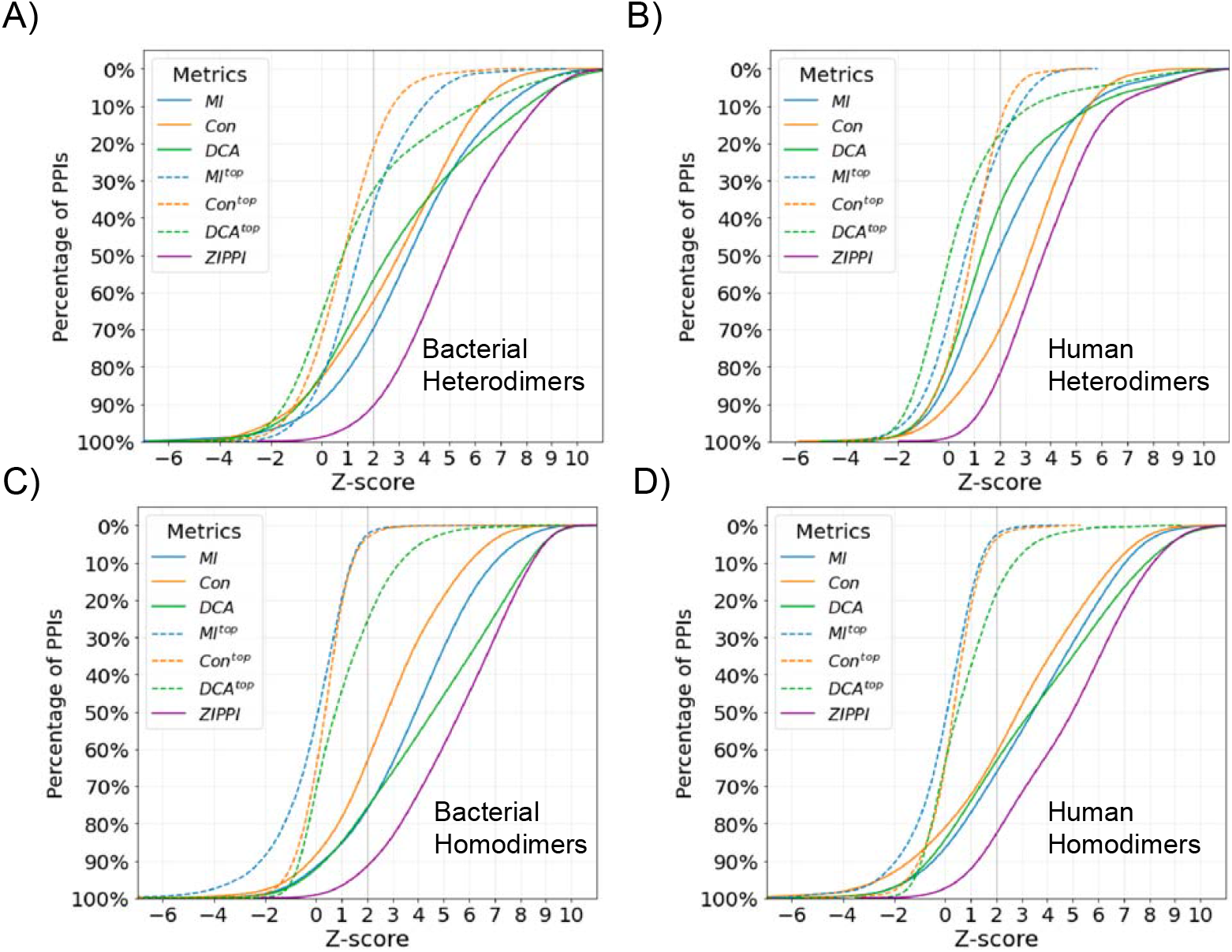
Percentage of PDB PPIs with a Z-score above a threshold. Colors indicate curves for different metrics each of which corresponds to the maximum of the raw and APC value for a given PPI. The mean and top metric of all interface contacts are denoted as <>, and ^top^, respectively The ZIPPI curve is shown in purple and, for a given PPI, is the largest Z-score among all metrics. A). Bacterial PDB heterodimers. B). Human PDB heterodimers. C). Bacterial PDB homodimers. D). Human PDB homodimers.

#### Heterodimers

The best performing metric for bacterial heterodimers is MI, followed by DCA (Figure 2A). About 90% of the PPIs have a ZIPPI-score > 2, ∼80% have a score > 3 and 65% have a score > 4. In contrast to bacteria, for human heterodimers the most significant metric is conservation (Figure 2B). We suggest that the difference is due to greater coevolutionary divergence underlying bacterial MSAs as opposed to eukaryotic MSAs. For human heterodimers about 82% of the PPIs have a ZIPPI-score > 2 and ∼63% have a ZIPPI-score > 3.

#### Homodimers

DCA is the best performing metric for both bacterial and human homodimers (Figures 2C,D), likely because for homodimers MI signal is strongest for coevolutionary coupling of intramolecular interactions rather versus intermolecular coupling. Secondly, the sequence depth in MSA is much larger (reflecting two copies of a single protein) than for paired alignments in heterodimer case, and DCA is thus more reliable for homodimers. The percent of bacterial homodimers that have a ZIPPI-score > 2, 3, 4 is ∼92%, 83% and 74% respectively. These values for human homodimers are ∼82%, 75% and 65%, respectively.

Overall, our results show that DCA works well for homodimers but that MI works best for heterodimers. This is likely a consequence of our analysis of interfacial residues alone and of the fact that we only consider observed contacts in a 3D structure.

#### Effect of MSA depth

Figure 3 plots ZIPPI score versus N_MSA_, the sequence depth of the MSAs. The figure also contains a histogram that displays the number of interfaces as a function of ZIPPI score. Surprisingly, there are cases where ZIPPI scores > 2 are obtained for values of N_MSA_ < 10. Most of these cases result from significant sequence conservation of interfacial residues but there are cases where even MI yields a significant signal. Although these few cases may well be statistical anomalies, there are many more interfaces with values of N_MSA_ between 10-50 where high Z-scores are calculated. These results highlight the success of ZIPPI in leveraging even shallow MSAs, made possible by the evaluation of interfacial residues in an experimentally determined structure. The next sections explore ZIPPI performance on predicted protein-protein complexes.

**Figure 3.**
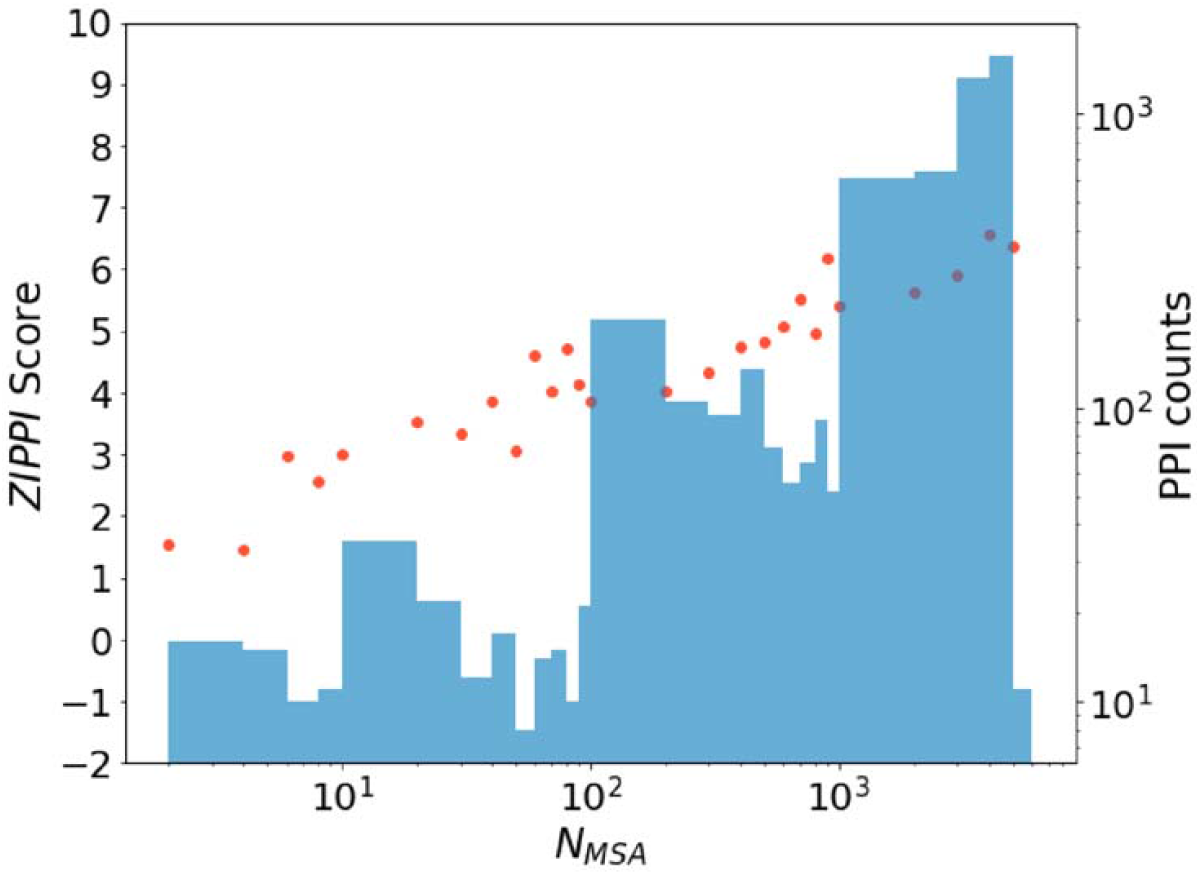
Effect of MSA depth on ZIPPI score for PDB dimers. The ZIPPI score is plotted against MSA depth, N_MSA_, where each red dot corresponds to the average ZIPPI score in a given bin. A histogram of the numbers of PPIs in each bin is shown in blue. Data is plotted on a log scale.

### Test on CAPRI benchmark decoys

We applied ZIPPI to a widely used decoy set, score_set^33^, to test CAPRI (Critical Assessment of Prediction of Interactions) models. This set contains docking models predicted by 47 different groups based on targets including proteins from bacteria, yeast, vertebrates and artificial design. We considered 13 widely studied targets for which there are 18,538 decoys, about 10% of which represent docking predictions of acceptable, medium or high quality based on CAPRI-defined criteria (we define this combined group as “acceptable+”) whereas the remaining are considered to be “incorrect.” Even though two of the targets, T53 and T54 contain designed proteins (both in T53 and one in T54),we found 2110 shared species for Target 53 and 198 shared species for Target 54. Table S1 reports MSA depth for all targets along with the number of acceptable+ and incorrect decoys, and the area under ROC curve (AUROC) for each target. It is clear from the table that extremely shallow MSA depths can produce good AUROCs and, further, that ZIPPI can handle cases with few interfacial residues and when there are only a few acceptable targets.

Figure 4 plots the percentage of all models that have a given ZIPPI score in each category across targets. There is a clear distinction between acceptable+ and incorrect decoys. Essentially 90% of the acceptable+ models have Z-scores > 2. Nevertheless, some incorrect decoys do have high Z-scores and some correct decoys have low Z-scores. The first of the two peaks in the high-quality curve is due to T40 that involves a trimeric complex between a bovine protein and two copies of the same plant protein which bind in different locations. Only one is considered in the decoy set but the other forms a second interface complicating the creation of non-interacting residues in randomly generated interfaces. This issue that does not affect docking approaches but compromises ZIPPI analysis.

**Figure 4.**
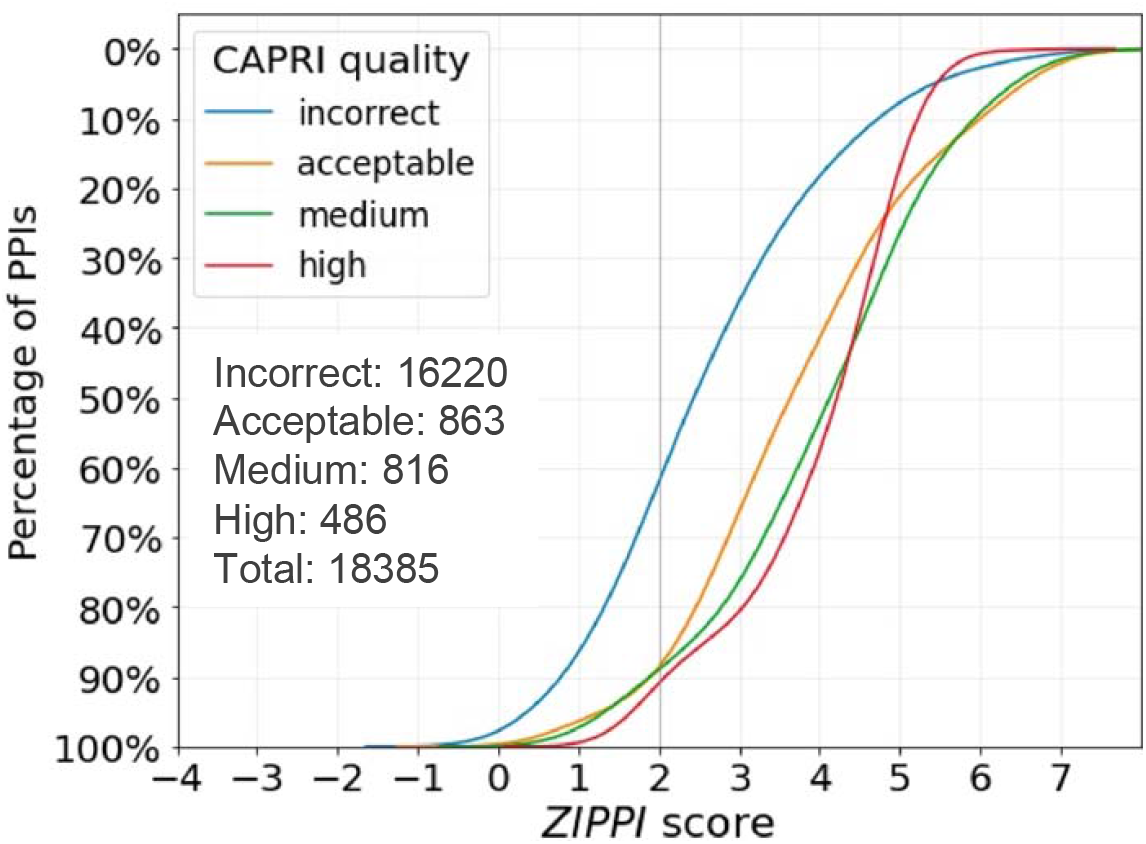
Percentage of CAPRI models having a given ZIPPI score. Percentages are plotted along the y-axis for four classes of CAPRI models. The total number of models in each class is indicated in the text at the lower left.

The AUROC and the Success Rate (the number of targets out of the total number considered for which at least one decoy of acceptable+ quality is present in the top N predictions), are widely used to evaluate CAPRI scoring functions. Table S2 lists AUROC and Success Rates for individual metrics as well as for the ZIPPI score. DCA, which is time consuming and not effective for heterodimers (Figure 2, Figure S1), is only used for homodimeric complexes. In contrast to the analysis of PDB complexes, all metrics perform approximately the same. Further, the improved performance of “top” relative to PDB complexes is likely because when not all contacts in an interface are real, incorrect contacts will lower the mean while only one correct contact needs to be predicted to yield a good “top” score. Of note, for this decoy set, an individual metric can produce a better score than the ZIPPI score (see Table S1A). This can result if some metric, e.g. Con, strongly favors an incorrect decoy and thus lowers the ZIPPI score for an acceptable+ model, while another metric, e.g. MI, favors an acceptable+ model. This can result in ZIPPI producing a false positive while MI chooses the acceptable+ model.

Table 1 compares ZIPPI performance to that of other methods^35–40^, most of which are based on machine learning. The data for other methods was taken from Table S8 of Réau *et al*.^36^ (see also Methods). ZIPPI, despite not being based on training, is essentially tied as the top performer as measured by AUROC and is the best performer based on top 100 Success Rate. However, ZIPPI is outperformed by a number of other methods as measured by top 1 and top 5 success rates while, based on these criteria, iScore is the best performer. Of note, AUROC is affected by the distribution of false and true positives in a list of predictions while Success Rate depends on the number of good predictions at the top of the list. Success Rates are central to CAPRI rankings while ROC curve performance may be more important in asking whether a particular prediction is correct.

**Table 1.** Performance of different scoring methods on CAPRI decoys. Area under the receiver operating characteristic curve. (AUROC) is calculated by averaging over values for each of 13 targets. Success Rates of Top N indicates the number of targets where there are acceptable or better predictions in the Top N predictions.

| Method | AUROC | Success Rates |  |  |
| --- | --- | --- | --- | --- |
|  |  | Top1 | Top5 | Top100 |
| ZIPPI | $0.72 \pm 0.13$ | 2/13 | 2/13 | 12/13 |
| HADDOCK | $0.57 \pm 0.23$ | 2/13 | 3/13 | 9/13 |
| iScore | $0.68 \pm 0.21$ | 5/13 | 6/13 | 9/13 |
| DeepRank | $0.64 \pm 0.19$ | 1/13 | 1/13 | 9/13 |
| DOVE | $0.56 \pm 0.14$ | 1/13 | 2/13 | 10/13 |
| GNN-DOVE | $0.63 \pm 0.16$ | 1/13 | 6/13 | 8/13 |
| DeepRank-GNN | $0.72 \pm 0.19$ | 1/13 | 5/13 | 10/13 |

### Relating ZIPPI scores to PrePPI modeling scores

In recent work we reported PrePPI calculations for the *E. coli* interactome focusing solely on the structural modeling score, SM, for interactions between structured domains^31^. The score was trained on the human HINT high-quality literature-curated (HINT-HQ-LC) dataset which is designed to contain high confidence binary interactions^41^. In that study, we analyzed the set of ∼9 million possible *E. coli* PPIs and were able to construct ∼5.4 million PrePPI models of varying quality. A ROC curve was reported for testing these models on the *E. coli* HINT-HQ-LC data set yielding an AUROC of 0.88, thus, attesting to the overall high-quality of the predictions.

Figure 5 displays whisker plots for the range of ZIPPI scores for PrePPI predictions in different FPR bins. As is clear from the diagram, as the PrePPI predictions become more reliable (lower FPR), the median ZIPPI score increases. These results provide a strong consistency check in that better structural models as defined by PrePPI produce stronger evolutionary signals as measured by ZIPPI. For heterodimers, at FPR < 10^−4^, the percentage of predicted PPIs with a ZIPPI score > 2, 3, 4 is 94%, 81% and 67%, respectively. The comparable numbers for PDB structures (see discussion of Figure 2) are 95%, 85% and 71% suggesting that PrePPI’s highest confidence predictions have ZIPPI scores close to those of PDB structures. Performance deteriorates as FPR increases but there are still many good ZIPPI scores for higher FPR values. Figure 5 demonstrates that high and low ZIPPI scores are obtained in all FPR bins suggesting ZIPPI score can be used as an additional evidence source for prioritizing PrePPI models.

**Figure 5.**
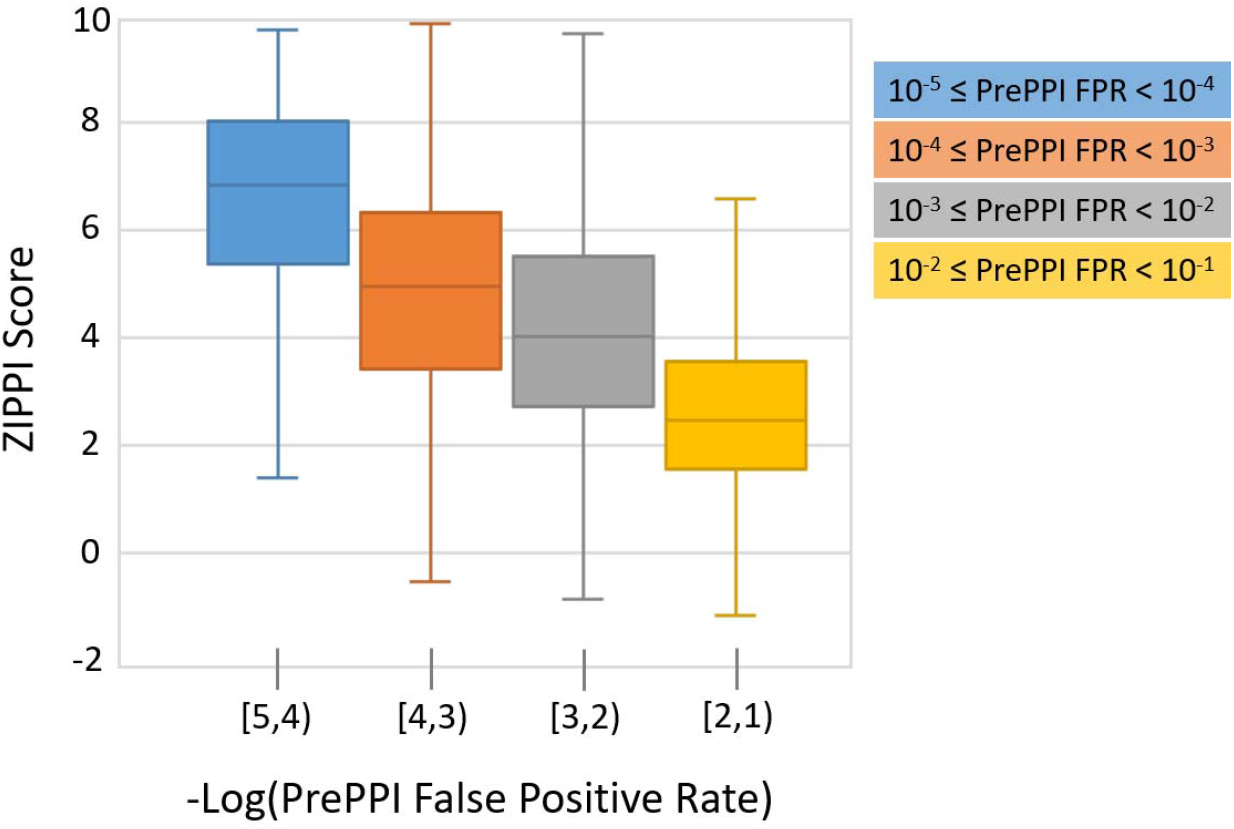
Relationship between ZIPPI scores and the False Positive Rate (FPR) of PrePPI-predicted PPIs. FPR ranges indicated below each set of bar charts and color-coded.

### The *E. coli* structural interactome

The size of the *E. coli* binary interactome has been estimated to be on the order of 10,000 but such estimates are necessarily, at the best, very rough approximations. Table 2 lists the number of proteins and number of PPIs (out of the 5.4 million predicted) for different FPR rates and different ZIPPI scores. At FPR < 0.01 PrePPI predicts ∼63,000 PPIs involving ∼3,500 proteins but these numbers are significantly decreased when more stringent PrePPI FPRs and ZIPPI scores are applied. Only ∼2,100 PPIs satisfy the restrictive criteria of FPR < 0.00001 and ZIPPI score > 4.

**Table 2.**
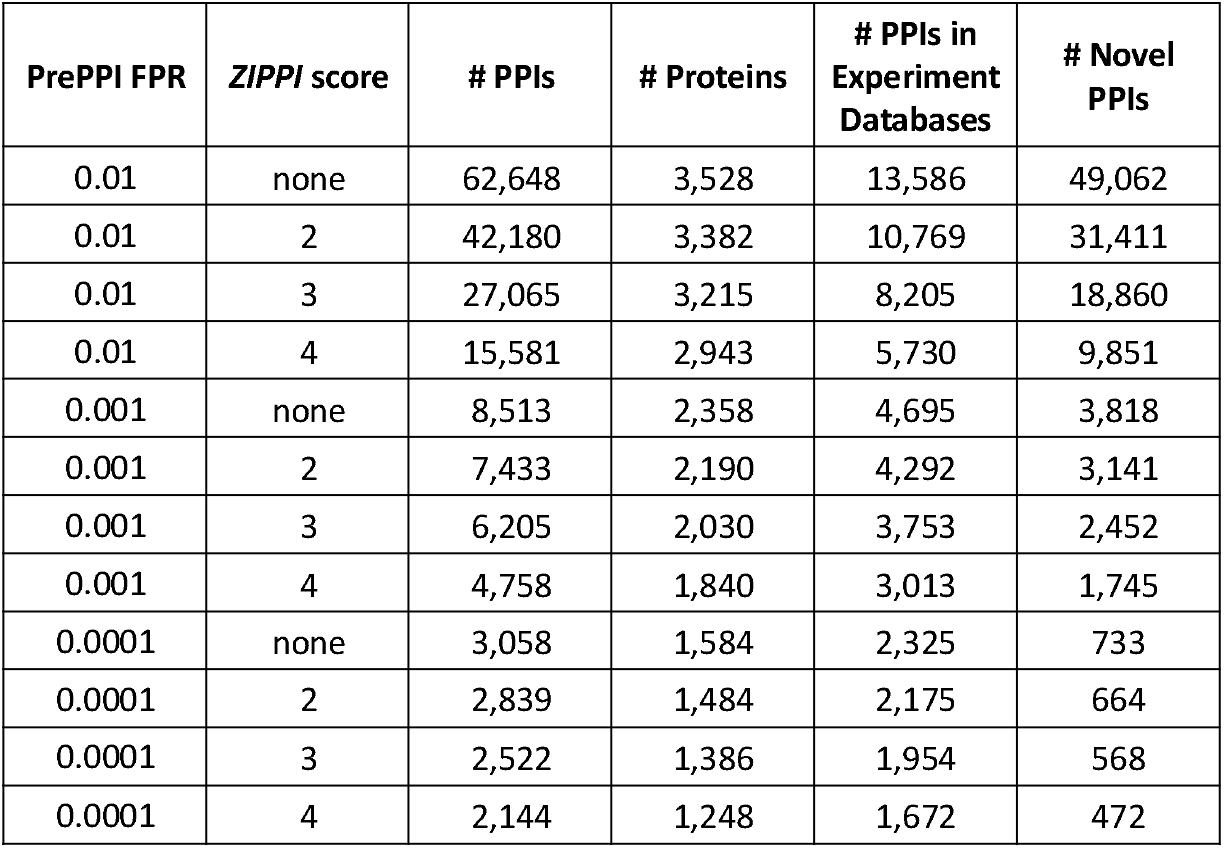
Number of proteins, PPIs and novel predictions for different PrePPI FPRs and different ZIPPI-scores.

Table 2 also lists the overlap of ZIPPI-filtered PrePPI predictions with PPIs annotated in experimental databases. Any PPI that appears in the listed databases (see methods) is considered whether or not the interaction is likely to be direct or indirect since our goal is to determine the number of truly novel PPIs that our methods predict. At the most stringent end of the scale (FPR < 0.0001, ZIPPI score > 4) 472 novel predictions are made. On the other hand, as an example, there are 9,851 predictions made for FPR < 0.01, ZIPPI score > 4 suggesting that using ZIPPI may facilitate the discovery of meaningful predictions that might be missed based on PrePPI alone (see Figure 5). In future work we plan to train classifiers based on both the PrePPI SM score and the ZIPPI score so as to more fully integrate the two methods.

## Discussion

Here we have introduced ZIPPI, a novel method that uses paired MSAs as a basis for scoring predicted models of protein-protein interfaces. ZIPPI leverages evolutionary information involving contacting residues in a 3D structural model rather than full length protein sequences. In addition to reducing the amount of computer time required to evaluate an interface, the fact that interacting residues are known from the structure obviates the need to carry out DCA and related methods to avoid the problem of indirect couplings. Methods such as AlphaFold-multimer^21^ and AF2Complex^22^ produce models of complexes but, due to computational limitations, have not been applied on an interactome-wide scale to ask which of all possible PPI pairs in a proteome will form a binary complex. However, in contrast, PrePPI uses structure to accomplish this task for the ∼200 million possible PPIs in human and for the ∼9 million possible PPIs in *E. coli*. More recently, the Threpp threading algorithm has also been used to screen the entire *E. coli* interactome for plausible structural models of complexes^42^. These studies highlight the fact that structure-based screening of entire interactomes is far more efficient than using sequence based deep learning structure-prediction methods for screening purposes. However, deep-learning based methods are in many cases likely to produce more accurate models than high-throughput computational procedures such as the ones used in PrePPI and Threpp. This suggests using structure-based approaches to provide interactome-wide yes/no answers along with 3D models and turning to increasingly accurate deep learning methods for a more limited set of interactions of particular interest. We emphasize that ZIPPI can quickly be applied to any structural model and is computationally efficient.

We first tested ZIPPI on bacterial and human complexes in the PDB and found that mutual information was the best scoring metric for bacterial heterodimers while sequence conservation was the most effective metric for human heterodimers. We found that DCA is not necessary for the evaluation of models of heterodimeric complexes when the interfacial contacts of a model are known and, rather, that the use of direct mutual information is sufficient. Moreover, given a structural model for a complex, we find that shallow MSAs are able to produce significant mutual information and conservation signals. DCA was found to be the most effective metric for homodimers and is retained in ZIPPI only for these cases.

ZIPPI was also tested on thirteen CAPRI targets which contained what are termed acceptable, medium and high quality models as well as a large number of incorrect decoys. Overall, ZIPPI performance was found to be comparable to or better than other approaches. Thus ZIPPI, especially if combined with other sources of evidence, may prove to be an effective means of evaluating future CAPRI predictions. We note in this regard that evolutionary information has been used for some time in the evaluation of docking models but usually in combination with other evidence sources. Nevretheless, ZIPPI’s focus on the use of MSAs to evaluate interfaces and its method of calculating Z-scores through the generation of random interfaces interfaces, offers a novel, computationally efficient and highly effective measure of interface quality that can easily be combined with other sources of evidence.

Finally, we implemented ZIPPI for 4.5 million *E. coli* PPI interfaces predicted by PrePPI. As suggested by the results in Table 2, the inclusion of ZIPPI in PrePPI predictions has the potential both to increase the reliability of “high confidence “predictions while identifying low confidence predictions that are worthy of further consideration. Despite its general applicability, an immediate application of ZIPPI is its combination with the PrePPI algorithm with the goal of combining evolutionary signals with a method based entirely on 3D structure. Indeed despite oversall correlation (Figure 5) the existence of PrePPI-predicted PPIs with high ZIPPI scores and low PrePPI-predicted FPRs (Table 2), indicates that the information in ZIPPI is highly complementary. Integration of the two methods should prove to be quite valuable, especially in applications to the human proteome and other eukaryotic organisms.

The combination of PrePPI with ZIPPI alters the prevailing paradigm of deep-learning-based approaches exemplified by AlphaFold-multimer which predict the structure of a complex from sequence alone. Rather, the strategy proposed here is to start with monomer structures (for example taken from AlphaFold as now done in PrePPI) and then to generate complex structures either through template-based modeling, as in PrePPI or, in principle, from docking. Our own focus is predicting PPIs on a proteome-wide scale as made possible with the PrePPI algorithm. ZIPPI now allows us to couple 3D structural information as provided by PrePPI with evolutionary information from MSAs. As such it opens the door to the integration of PrePPI, not only with ZIPPI but also with deep learning based approaches that are computationally efficient enough to provide residue-contact probabilities on a proteome-wide scale which for human, may involve billions of PPIs if predictions are to be made at the domain level.

## Methods

### Selecting bacterial and human PDB dimer structures

Taxonomy and UniProtKB summary files for all PDB chains were downloaded from SIFTS^43^. From the SIFTS PDB chain taxonomy file, PDB chains that only correspond to one taxonomy ID were selected and then filtered to bacteria and human PDB chains. The taxonomy list of bacteria was collected from searching both UniProtKB-proteome^44^ and NCBI Taxonomy databases^45^. The union of the two databases provided 521,897 unique bacteria taxonomy IDs.

From the SIFTS PDB chain UniProt file, PDB files with only two UniProt IDs for heterodimers and one ID for homodimers, two chains and both chains longer than 30 amino acids are selected. PDBs that have any chain mapped to ≥ 2 UniProt IDs are excluded to avoid fusion or chimera proteins. Structure resolution information is obtained through the PDB API service^27^. PDBs that are protein-only as the polymer entity type, and either from X-ray with resolution ≤ 4 Å or from EM with resolution ≤ 4.5 Å are selected. NMR structures are not used. Further, through reading the PDB file header, PDBs where the oligomer state of the first BioAssembly (*aka*. BioUnit or BioMolecule) defined as “DIMERIC” by either the author or software with resolved sequence lengths longer than 30 amino acids are selected. PDB dimer structures that have redundant UniProt ID pairs are removed by keeping the structure with either longer length (at least twice as long) or better structural resolution. Lastly, to remove closely related homolog proteins, we compared the pairwise sequence identities and removed redundant structures where both protein sequences have 90% sequence identity with another structure. The detailed pipeline is provided in the supplemental information.

### Defining protein surface and protein-protein interface

The accessible surface area (ASA) of residues for individual chain A, B, and their complex AB are obtained using our in-house C++ program AREA. An interface is defined as long as the buried ASA larger than zero. The interface between proteins A and B consists of contacting residues where the distance between any heavy atoms is less than 6.0 Å. All the residue indices from the PDB are updated after mapping the PDB sequences to their full UniProt sequences using hhalign of the hh-suite package^46^.

### Generating random protein-protein interfaces

The interfacial residues on proteins A and B are replaced, one by one, by randomly chosen surface residues of the same protein as indicated in Figure 1. This way of generating random interfaces preserves the interaction network of different interfacial residues. To ensure statistical significance of the Z-score calculations, 100 random interfaces are generated for each protein-protein interface.

### Generating and pairing MSAs

To avoid biased sequence sampling due to over-studied model species, we carried out homolog sequence search on 5,090 representative proteomes that were carefully curated and selected in EggNog 5.0^47^. This database includes 4,445 prokaryotic reference genomes selected from original 25,038 bacteria genomes, and 477 eukaryotic genomes. Homologous sequences are searched using Jackhmmer (hmmer-3.2.1)^48^ with 5 iterations and the default E-value of 0.001. In the final outputted multiple sequence alignment, only the sequence with highest identity to the query is kept as the representative sequence for each species.

The MSAs of two proteins, p1 and p2, are paired based on the shared common species. Sequence rows that cover less than 50% of surface residue positions of p1 or p2 are excluded from the paired MSA. Sequence columns, either interface residual or surface residual positions, that have more than 50% gaps are excluded.

### Calculating mutual information, conservation, DCA, and their APC-corrected terms

For two positions (***a, b***) in the paired MSA, their mutual information (MI) is calculated through Eq. 1, where *x* and *y* denote their amino acid type and the gap in MSA is treated as the 21th amino acid type or state. The *p(x)* and *p(y)* are the frequency of a certain amino acid types and *p(x, y)* is the frequency of a certain pairs of amino acid types. The conservation score between two positions (***a, b***) is defined through the complement of their normalized joint entropy *S(a, b)* (Eq. 2). The direct coupling information is calculated through the mean field DCA method which is based on the maximum-entropy model^3^. The final direct coupling information is quantified using a similar definition as in the mutual information except *p*^*(dir)*^*(x, y)* involves only the isolated direct coupling strength of (***a, b***) from the DCA calculations (Eq. 3).

The average product correction (APC) is applied to all measurements. Taking MI as an example, the APC term between position (***a, b***) (from p1 and p2, respectively) is calculated as the product of the average MI signal of position ***a*** with positions of p2, and position ***b*** with positions of p1, between interfacial residues on both proteins, then normalized by the average measurement of all protein to the other (Eq. 4). The APC-corrected term is given correspondingly for MI, Con, and DCA (Eq. 5).

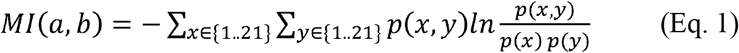

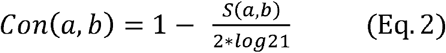

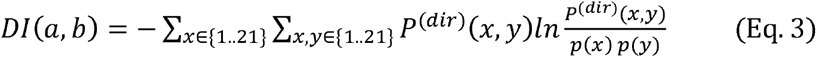

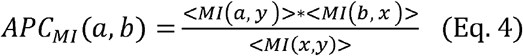

### Calculating Z-scores of the interface

For each interface contact of the given interface between protein p1 and p2 and the generated 100 random interfaces, the following six measurements are calculated: mutual information, conservation, direct coupling information and their corresponding APC-corrected terms. Of all the interface contacts, we choose the top and the mean as the representative metric for each measurement, denoted as MI^top^ and <MI>, for example. The Z-score of the 12 metrics are then calculated for the given interface versus the generated random interfaces. The larger Z-score of the raw metric and its APC-corrected metric is taken as Z-score for this metric. The maximum of all metrics is taken as the final ZIPPI score.

### Building the *E. coli* experimental PPI database

The experimental database of *E. coli* PPI is integrated from several major resources including Interactome3D^49^, HINT^41^, APID^50^, STRING^51^ and Ecocyc^52^, as well as previously known large-scale *E. coli* PPI screening using experimental methods such as APMS^53^ and Y2H^54^. Before their integration, each database was pre-processed by selecting only *E. coli* K12 proteins (proteome size: 4391) and sorting the uniport IDs for each pair of PPIs. During the integration, redundant PPIs were removed. Note that Interactome3D also includes homology-modeled PPIs and the STRING database has inferred PPIs, which are not determined by direct physical interaction experiments but inferred by other methods such as gene-related methods or species PPI transfer. By excluding these two contributions, we also built a purely experimental PPI database of *E. coli* based on direct physical experiments. In all, there are 565,007 PPIs in the integrated experimental database set and 45,634 PPIs in the physical experimental PPI dataset.

In summary, the integrated experimental database set includes: all HINT binary and complex PPIs (updates of 2021/11), all APID PPIs (updates of 2021/11), all Interactome3D PPIs (updates of 2021/11), all STRING PPIs (v11.5), the gold standard dataset used in Zhang and coworkers’ Threpp work^42^, the high throughput experimental PPI set from Threpp (Table S1), the Ecocyc PPIs downloaded Cong *et al*.^55^(Table S5), the Y2H PPI set from Rajagopala *et al*.^54^ (Supplementary Table 2), the high-confidence and median-confidence APMS PPI set from Babu *et al*.^53^ (Supplementary Table 2). For the physical experimental PPI dataset, only physical links in the STRING database with experimental score >0 are included; only the PDB subset of Interactome3D is included; the other datasets remain the same as in the integrated experimental database.

---

**SI Figure S1.**
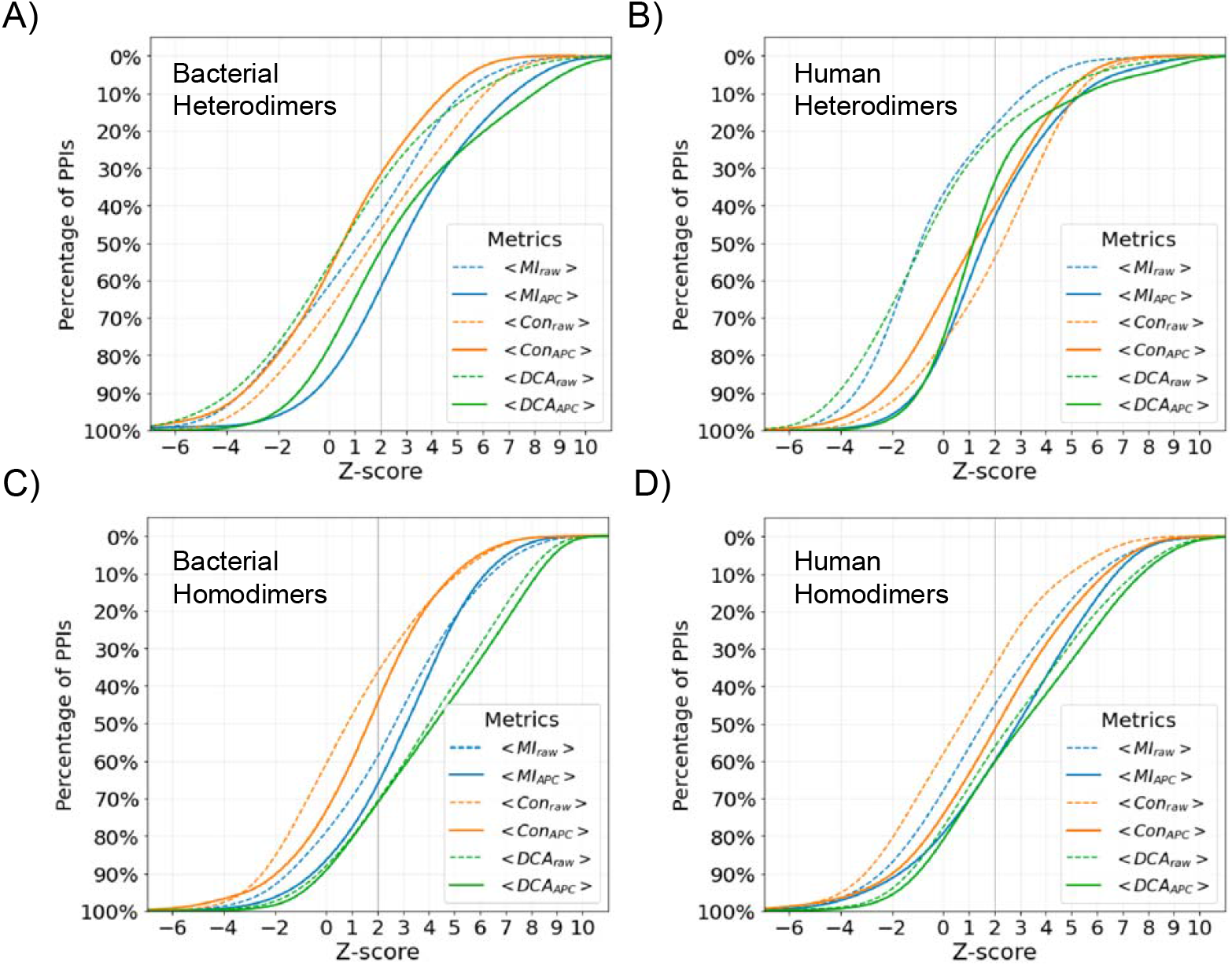
Percentage of PDB PPIs with a Z-score above a threshold for the raw and APC-corrected metrics averaged over interface contacts. Colors indicate curves for different metrics A). Bacterial PDB heterodimers. B). Human PDB heterodimers. C). Bacterial PDB homodimers. D). Human PDB homodimers.

**SI Figure S2.**
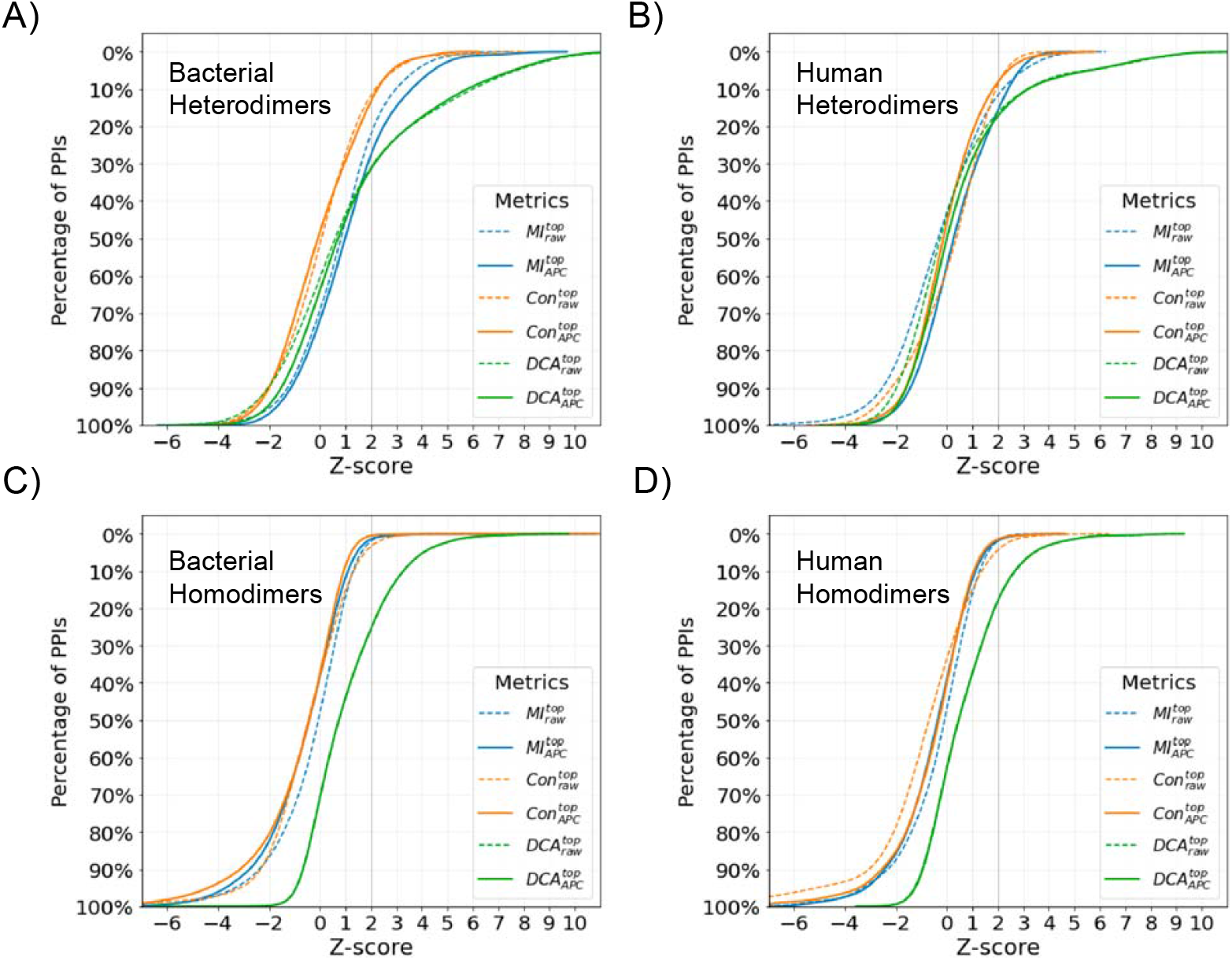
Percentage of PDB PPIs with a Z-score above a threshold for the interface contact with the top value for a given metric. Colors indicate curves for different metrics. A). Bacterial PDB heterodimers. B). Human PDB heterodimers. C). Bacterial PDB homodimers. D).Human PDB homodimers.

**SI Table S1.** Properties associated with each CAPRI target.

| ID | $N_{MSA}$ | $< N_{IFR} >$ | $N_{\geq accpt.}$ | $N_{incorrect}$ | AUROC (ZIPPI) |
| --- | --- | --- | --- | --- | --- |
| Target29 | 623 | 64 | 167 | 1916 | 0.926 |
| Target30 | 155 | 42 | 2 | 1341 | 0.550 |
| Target32 | 62 | 70 | 15 | 584 | 0.687 |
| Target35 | 161 | 65 | 3 | 496 | 0.528 |
| Target37 | 142 | 48 | 99 | 1401 | 0.542 |
| Target39 | 267 | 58 | 4 | 1396 | 0.843 |
| Target40 | 55 | 61 | 588 | 1592 | 0.664 |
| Target41 | 27 | 50 | 371 | 829 | 0.755 |
| Target46 | 1396 | 56 | 24 | 1675 | 0.651 |
| Target47 | 24 | 53 | 611 | 440 | 0.930 |
| Target50 | 88 | 3 | 133 | 1318 | 0.734 |
| Target53 | 2112 | 55 | 130 | 1270 | 0.742 |
| Target54 | 198 | 49 | 19 | 1381 | 0.763 |

**SI Table S2.**
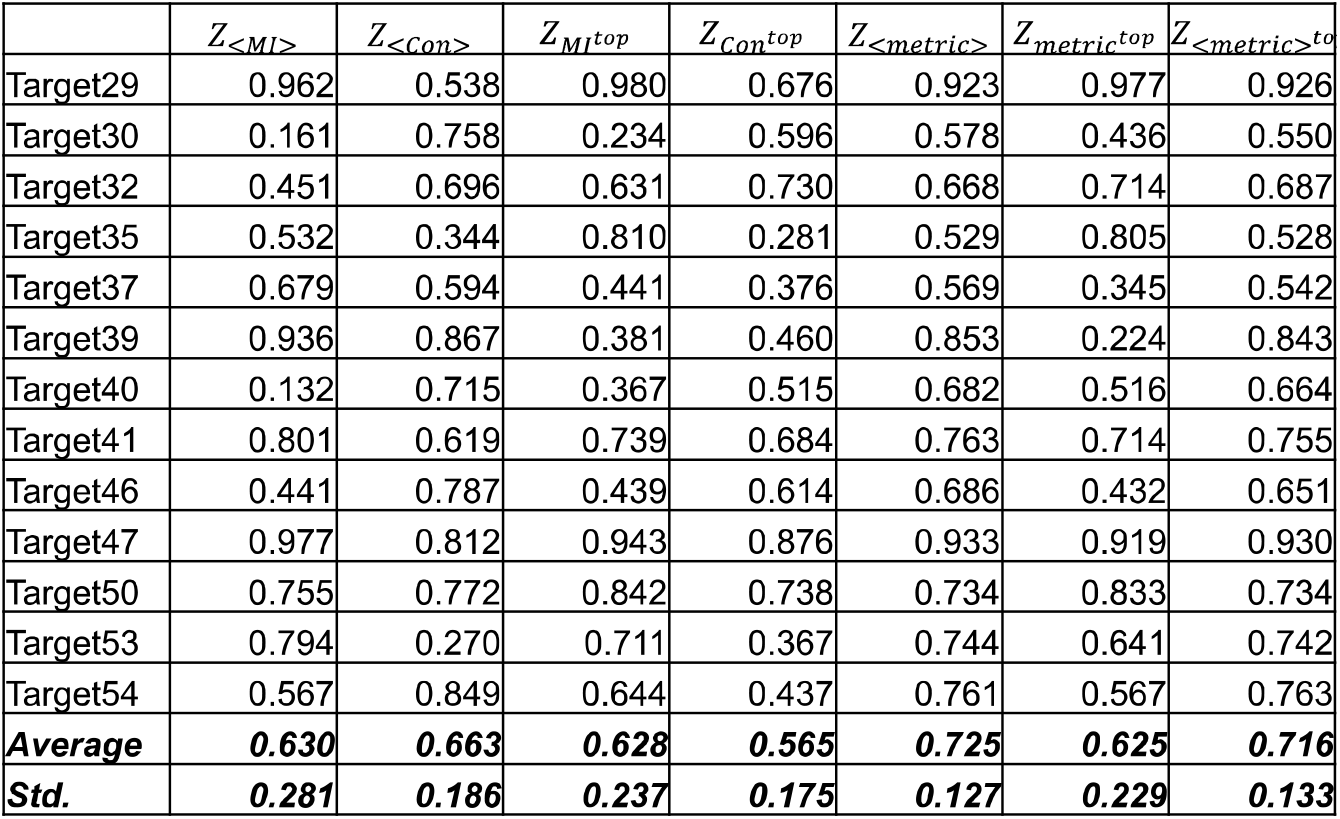
Detailed AUROC performances of ZIPPI metrics for each CAPRI Target.

## Notes

### Competing Interest Statement

The authors have declared no competing interest.

## References

1. Göbel, U., Sander, C., Schneider, R. & Valencia, A. Correlated mutations and residue contacts in proteins. Proteins: Structure, Function, and Bioinformatics 18, 309–317 (1994).

2. Shoemaker, B. A. & Panchenko, A. R. Deciphering protein-protein interactions. Part II. Computational methods to predict protein and domain interaction partners. PLoS Computational Biology vol. 3 595–601 Preprint at https://doi.org/10.1371/journal.pcbi.0030043 (2007).

3. Morcos, F. et al. Direct-coupling analysis of residue coevolution captures native contacts across many protein families. Proc Natl Acad Sci U S A (2011) doi:10.1073/pnas.1111471108.

4. Kamisetty, H., Ovchinnikov, S. & Baker, D. Assessing the utility of coevolution-based residue-residue contact predictions in a sequence- and structure-rich era (Proceedings of the National Academy of Sciences of the United States of America (2013) 110, 39 (15674-15679) DOI:10.1073/pnas.1314045110). Proc Natl Acad Sci U S A 110, 18734–18735 (2013).

5. Marks, D. S. et al. Protein 3D structure computed from evolutionary sequence variation. PLoS One 6, (2011).

6. Hopf, T. A. et al. Sequence co-evolution gives 3D contacts and structures of protein complexes. Elife 3, 1–45 (2014).

7. Ovchinnikov, S., Kamisetty, H. & Baker, D. Robust and accurate prediction of residue-residue interactions across protein interfaces using evolutionary information. Elife 2014, 1–21 (2014).

8. Green, A. G. et al. Large-scale discovery of protein interactions at residue resolution using co-evolution calculated from genomic sequences. Nat Commun 1–12 (2021) doi:10.1038/s41467-021-21636-z.

9. Bitbol, A. F., Dwyer, R. S., Colwell, L. J. & Wingreen, N. S. Inferring interaction partners from protein sequences. Proc Natl Acad Sci U S A 113, 12180–12185 (2016).

10. Levy, R. M., Haldane, A. & Flynn, W. F. Potts Hamiltonian models of protein co-variation, free energy landscapes, and evolutionary fitness. Current Opinion in Structural Biology vol. 43 55–62 Preprint at https://doi.org/10.1016/j.sbi.2016.11.004 (2017).

11. Ekeberg, M., Lövkvist, C., Lan, Y., Weigt, M. & Aurell, E. Improved contact prediction in proteins: Using pseudolikelihoods to infer Potts models. Phys Rev E Stat Nonlin Soft Matter Phys 87, (2013).

12. Weigt, M., White, R. A., Szurmant, H., Hoch, J. A. & Hwa, T. Identification of direct residue contacts in protein-protein interaction by message passing. Proc Natl Acad Sci U S A 106, 67–72 (2009).

13. Jones, D. T., Buchan, D. W. A., Cozzetto, D. & Pontil, M. PSICOV: Precise structural contact prediction using sparse inverse covariance estimation on large multiple sequence alignments. Bioinformatics 28, 184–190 (2012).

14. Jumper, J. et al. Highly accurate protein structure prediction with AlphaFold. Nature 596, 583–589 (2021).

15. Baek, M. et al. Accurate prediction of protein structures and interactions using a 3-track network. Science (1979) 8754, 2021.06.14.448402 (2021).

16. Wang, S., Sun, S., Li, Z., Zhang, R. & Xu, J. Accurate De Novo Prediction of Protein Contact Map by Ultra-Deep Learning Model. PLoS Computational Biology vol. 13 (2017).

17. Aq2qaaq21234567890-Bepler, T. & Berger, B. Learning the protein language: Evolution, structure, and function. Cell Syst 12, 654–669.e3 (2021).

18. Sledzieski, S., Singh, R., Cowen, L. & Berger, B. D-SCRIPT translates genome to phenome with sequence-based, structure-aware, genome-scale predictions of protein-protein interactions. Cell Syst 12, 969–982.e6 (2021).

19. Lin, Z. et al. Evolutionary-scale prediction of atomic-level protein structure with a language model. Science vol. 379 https://www.science.org (2023).

20. Chowdhury, R. et al. Single-sequence protein structure prediction using a language model and deep learning. Nat Biotechnol 40, 1617–1623 (2022).

21. Evans, R. et al. Protein complex prediction with AlphaFold-Multimer. (2022) doi:10.1101/2021.10.04.463034.

22. Gao, M., Nakajima An, D., Parks, J. M. & Skolnick, J. AF2Complex predicts direct physical interactions in multimeric proteins with deep learning. Nat Commun 13, (2022).

23. Humphreys, I. et al. Computed structures of core eukaryotic protein complexes. Science (1979) 374, (2021).

24. Vakser, I. A. Protein-protein docking: From interaction to interactome. Biophysical Journal vol. 107 1785–1793 Preprint at https://doi.org/10.1016/j.bpj.2014.08.033 (2014).

25. Barradas-Bautista, D., Rosell, M., Pallara, C. & Fernández-Recio, J. Structural Prediction of Protein–Protein Interactions by Docking: Application to Biomedical Problems. in Advances in Protein Chemistry and Structural Biology vol. 110 203–249 (Academic Press Inc., 2018).

26. Petrey, D. et al. Template-based prediction of protein function. Current Opinion in Structural Biology vol. 32 33–38 Preprint at https://doi.org/10.1016/j.sbi.2015.01.007 (2015).

27. Berman, H. M. et al. The Protein Data Bank. Nucleic Acids Research vol. 28 http://www.rcsb.org/pdb/status.html (2000).

28. Zhang, Q. C., Petrey, D., Norel, R. & Honig, B. H. Protein interface conservation across structure space. Proc Natl Acad Sci U S A 107, 10896–10901 (2010).

29. Zhang, Q. C. et al. Structure-based prediction of protein-protein interactions on a genome-wide scale. Nature 490, 556–560 (2012).

30. Garzón, J. I. et al. A computational interactome and functional annotation for the human proteome. Elife 5, 1–27 (2016).

31. Petrey, D., Zhao, H., Trudeau, S., Murray, D. & Honig, B. PrePPI: A structure informed proteome-wide database of protein-protein interactions. J Mol Biol 168052 (2023) doi:10.1016/j.jmb.2023.168052.

32. Janin, J. et al. CAPRI: A critical assessment of PRedicted interactions. Proteins: Structure, Function and Genetics 52, 2–9 (2003).

33. Lensink, M. F. & Wodak, S. J. Score_set: A CAPRI benchmark for scoring protein complexes. Proteins: Structure, Function and Bioinformatics 82, 3163–3169 (2014).

34. Dunn, S. D., Wahl, L. M. & Gloor, G. B. Mutual information without the influence of phylogeny or entropy dramatically improves residue contact prediction. Bioinformatics 24, 333–340 (2008).

35. Renaud, N. et al. DeepRank: a deep learning framework for data mining 3D protein-protein interfaces. Nat Commun 12, 1–8 (2021).

36. Réau, M., Renaud, N., Xue, L. C. & Bonvin, A. M. J. J. DeepRank-GNN: a graph neural network framework to learn patterns in protein-protein interfaces. Bioinformatics 39, 1–8 (2023).

37. Geng, C. et al. Structural bioinformatics iScore: a novel graph kernel-based function for scoring protein-protein docking models. Bioinformatics 36, 112–121 (2019).

38. Dominguez, C., Boelens, R. & Bonvin, A. M. J. J. HADDOCK: A protein-protein docking approach based on biochemical or biophysical information. J Am Chem Soc 125, 1731–1737 (2003).

39. Wang, X., Flannery, S. T. & Kihara, D. Protein Docking Model Evaluation by Graph Neural Networks. Front Mol Biosci 8, (2021).

40. Wang, X., Terashi, G., Christoffer, C. W., Zhu, M. & Kihara, D. Protein docking model evaluation by 3D deep convolutional neural networks. Bioinformatics 36, 2113–2118 (2020).

41. Das, J. & Yu, H. HINT: High-quality protein interactomes and their applications in understanding human disease. BMC Syst Biol 6, (2012).

42. Gong, W. et al. Integrating Multimeric Threading With High-throughput Experiments for Structural Interactome of Escherichia coli. J Mol Biol 433, 166944 (2021).

43. Velankar, S. et al. SIFTS: Structure Integration with Function, Taxonomy and Sequences resource. Nucleic Acids Res 41, (2013).

44. Bateman, A. et al. UniProt: the universal protein knowledgebase in 2021. Nucleic Acids Res 49, D480–D489 (2021).

45. Federhen, S. The NCBI Taxonomy database. Nucleic Acids Res 40, (2012).

46. Remmert, M., Biegert, A., Hauser, A. & Söding, J. HHblits: Lightning-fast iterative protein sequence searching by HMM-HMM alignment. Nat Methods 9, 173–175 (2012).

47. Huerta-Cepas, J. et al. EggNOG 5.0: A hierarchical, functionally and phylogenetically annotated orthology resource based on 5090 organisms and 2502 viruses. Nucleic Acids Res 47, D309–D314 (2019).

48. Eddy, S. R. HMMER User’s Guide Biological sequence analysis using profile hidden Markov models. http://hmmer.org (2020).

49. Mosca, R., Céol, A. & Aloy, P. Interactome3D: Adding structural details to protein networks. Nat Methods 10, 47–53 (2013).

50. Alonso-López, Di. et al. APID database: Redefining protein-protein interaction experimental evidences and binary interactomes. Database 2019, (2019).

51. Szklarczyk, D. et al. The STRING database in 2021: Customizable protein-protein networks, and functional characterization of user-uploaded gene/measurement sets. Nucleic Acids Res 49, D605–D612 (2021).

52. Keseler, I. M. et al. The EcoCyc database: Reflecting new knowledge about Escherichia coli K-12. Nucleic Acids Res 45, D543–D550 (2017).

53. Babu, M. et al. Global landscape of cell envelope protein complexes in Escherichia coli. Nat Biotechnol 36, 103–112 (2018).

54. Rajagopala, S. V. et al. The binary protein-protein interaction landscape of escherichia coli. Nat Biotechnol 32, 285–290 (2014).

55. Cong, Q., Anishchenko, I., Ovchinnikov, S. & Baker, D. Protein interaction networks revealed by proteome coevolution. Science (1979) 365, 185–189 (2019).

